# Chemotagging: a chemogenetic approach for identifying cell types with *in vivo* calcium imaging

**DOI:** 10.1101/2024.11.27.625756

**Authors:** BumJin Ko, Madeline E. Bacon, Yosif Zaki, Denise J. Cai

## Abstract

The ability to monitor the activity of specific cell types *in vivo* is critical for understanding the complex interplay between various neuronal populations driving freely moving behavior. Existing methods, such as optogenetic tagging (i.e., Optotagging^1^), have proven useful for identifying cell types in *in vivo* electrophysiological recordings during freely moving behavior. However, electrophysiological recordings are limited in their capacity to track the same neuronal populations across long periods of time (days to weeks). Single-photon miniscope imaging offers the advantage of tracking the same cells across weeks to months; however, it is difficult to distinguish different cell types within the recorded population. Here, we present “chemotagging,” a technique that allows for the identification of specific cell types in *in vivo* calcium imaging recordings. This protocol offers a method for tagging cell types with chemogenetic tools like Designer Receptors Exclusively Activated by Designer Drugs (DREADDs)^2^, while simultaneously recording calcium activity from a pan-neuronal population with calcium indicators. We highlight the key advantages and limitations of chemotagging and its potential implications for neuroscience research.

## Introduction

The brain is an intricate structure built from interconnections of cells. Advances in genomic profiling have significantly enhanced our understanding of the diverse cell types that form this network and their roles in encoding information^3^. Over the past few decades, research has revealed that the mammalian brain contains more than five thousand distinct cell-type clusters based on their gene expression^4^, each potentially serving unique functional roles^5,6^.

These distinct cell types exhibit heterogeneous patterns of activity. Moreover, their activity patterns are highly dynamic and influenced by multiple factors, including external stimuli^7^. To better understand how the brain operates across time and experience, it is crucial to track the same network of cells over time, while accurately distinguishing between these various cell types. The advent of calcium imaging techniques, such as two-photon microscopy^8^ and single-photon miniaturized fluorescence microscopes (miniscope)^9,10^, has revolutionized our ability to monitor large populations of neurons *in vivo*. These methods allow the real-time tracking of neuronal activity across weeks to months, providing invaluable insight into how networks of cells across different brain regions and neuronal circuits function. miniscopes enable imaging in animals when they are freely moving. However, they suffer from lower spatial resolution and shorter focal length compared to two-photon imaging^11^. GRIN (gradient index) lenses, essential for imaging deeper brain regions, are prone to chromatic aberration. Since these lenses bend light based on a gradient refractive index, different wavelengths (e.g., from multiple fluorophores or dual-channel imaging setups) often are refracted at different angles, leading to misalignment of images from different channels. One way to address this challenge is to use a recently developed dual-channel miniscope system that accounts for the chromatic aberration^12^, however, this requires specialized hardware.

One common approach to identify specific cell types during *in vivo* electrophysiological recordings is “optotagging,” which utilizes optogenetics to activate specific cell population for identification.^13^ However, optotagging has several key drawbacks when performed with miniscopes. One major issue is that red-shifted opsins, such as Chrimson^14^, often have long activation tails in shorter wavelengths of light, which overlaps with the excitation spectrum of GFP-based calcium indicators, such as GCaMP^15^, which interferes with the distinction between physiological and optogenetically driven signals. Switching from GFP-based calcium indicators to red-shifted ones also presents several challenges. Red-shifted calcium indicators, like RCaMP^16^, have lower brightness compared to GFP-based ones. This can result in a low signal-to-noise ratio which can mask subtle changes in neural activity. These factors make optotagging challenging with single-photon calcium imaging setup.

To enable the simultaneous imaging of multiple cell types with miniscopes, we propose a novel technique termed “chemotagging” that we recently published^17^. This method employs chemogenetic tools, such as DREADDs^2^, to selectively activate G-protein-coupled receptor (GPCR) signaling in specific cell types, while simultaneously using calcium imaging to monitor intracellular calcium ion dynamics^18-20^. Chemotagging enables the tagging of specific cell types without the drawbacks associated with optogenetics or using red-shifted calcium indicators. Here, we present an example of identifying inhibitory neurons among a large-scale pan-neuronal recording in the mouse hippocampus^17^. This technique is versatile and can be flexibly applied to different cell types for circuit-specific investigations. Moreover, this approach can be combined with both two-photon imaging or dual-channel miniscope setups, potentially offering greater flexibility and specificity in experimental design. This adaptability makes chemotagging a powerful tool for advancing the study of neural circuits across various experimental models.

### Procedures

1. **Planning** Choose a target cell type and an excitatory DREADD system. We used a *Gad2*-Cre mouse line and a Cre-dependent excitatory DREADD virus (rAAV5-DIO-hSyn-hM3Dq-mCherry), labeling inhibitory neurons that would later be activated by CNO. *Timing: Not Applicable*.
  - There are a number of other types of subpopulations that can be targeted using this DREADD system. For example, one could label genetically defined populations, as employed here, based on the expression of a particular gene. This could leverage AAVs with a promoter that targets a specific cell type of interest, or with a Cre mouse line that expresses Cre under the control of a specific promoter (as employed here). Similarly, one could target activity-dependent ensembles of neurons as well, labeling cells that express an immediate early gene during specified windows of time^21^. Alternatively, one could label projection-specific populations, for example employing a retrograde Cre-expressing virus in one brain area, and then injecting a Cre-dependent excitatory DREADD in an upstream brain area^2^. Here, we outline a protocol targeting inhibitory neurons using *Gad2*-Cre mice and a Cre-dependent excitatory DREADD virus. Please adjust the steps as necessary for whichever technique is best suited to target a given cell type.
2. **Viral infusions:** Decide which brain region will be targeted for calcium imaging based on the question being addressed in the given experiment. We performed our experiments in the dorsal CA1 of the hippocampus. Prepare a virus cocktail, mixing rAAV1-Syn-GCaMP6f with rAAV5-Syn-DIO-hM3Dq-mCherry in a 1:1 ratio. Infuse the appropriate volume of this virus cocktail into the desired brain region, as previously described^22^; we infused 300nL of virus to express in the pyramidal layer of dorsal CA1 of the hippocampus. *Timing: 20-60 minutes per mouse*.
  - Ensure that the viral titer is appropriate both for expression of the calcium indicator as well as of the hM3Dq receptor. This might have to be tested empirically for new lots of virus. Keep in mind that mixing the GCaMP and DREADD viruses will halve the titer of each virus. Consider performing a dilution study to decide on an optimal dilution, as previously described.^22^
3. **Lens implantation:** Wait two weeks for the mouse to recover after viral injection. Conduct a second surgery to implant a GRIN lens above the previously infused injection site, performing lens implantation as previously described^17^. Note that these two surgeries can also be performed together^22^. In this case, twenty minutes after the viral infusion is sufficient for virus diffusion. *Timing: 60-90 minutes per mouse*.
4. **Baseplating**: Allow another two weeks for the mouse to recover from surgery and for inflammation from the lens implantation to reduce. Conduct a final procedure to cement a baseplate on the mouse’s head above the GRIN lens, as previously described^22^. The miniscope will later attach to this baseplate to perform calcium imaging. This procedure is visually guided by the miniscope to find the optimal field-of-view which resolves the most neuronal cell bodies in the best focal plane. *Timing: 30-60 minutes per mouse*.
5. **Run calcium imaging experiment:** This step will vary for each use case, likely from days to months, depending on the experimental paradigm. In our experiment, we performed calcium imaging of dorsal CA1 neurons while mice were exposed to a novel environment paired with a footshock (Figure 2 Aversive). We then placed mice back into their homecage and recorded calcium dynamics while mice rested during this offline period (Figure 2 Offline). This experiment was designed to measure the offline reactivation of neuronal ensembles that were previously active during encoding. *Timing: variable depending on the experiment, one day in this case*.
6. **Chemotag the target cell type**: Choose an exogenous ligand to chemogenetically activate the hM3Dq receptors in the target cell type. We chose clozapine-N-oxide dihydrochloride (CNO) because it readily dissolves in saline and can easily be administered intraperitoneally. On the day after the experiment has ended, attach the miniscope to the mice, inject mice intraperitoneally with CNO at 3 mg/kg and place them back in their homecage. Immediately after the CNO injection, record calcium for the next 45 minutes to measure chemogenetically induced changes in calcium dynamics (Figure 2 Chemotag Session). This procedure can be conducted at any point after the conclusion of the calcium imaging experiment. We opted to chemotag one day after our final recording. *Timing: 60 minutes*.
  - The exogenous ligand and optimal dose that is employed will have to be empirically tested, based on the ligand and dose that maximally alters calcium dynamics after ligand administration *in vivo*.
7. **Pre-process calcium imaging data**: For each recording session, process the calcium recordings through a cell segmentation algorithm of choice. For this, we used Minian, an open-source interactive calcium imaging processing pipeline developed by our lab^23^. Perform this for each session and extract the spatial footprints, which represent the pixel locations that make up each recorded cell. In our experiment, we extracted the spatial footprints for cells recorded during Aversive encoding, during the subsequent Offline period, and during the Chemotag session on the following day. *Timing: Depends on length and number of recordings. In our case, one day*.
  - Users may alternatively use other calcium imaging processing pipelines (e.g., CaImAn^24^, MIN1PIPE^25^, etc).
8. **Cross-register cells across sessions**: Take the spatial footprints recorded across all recording sessions and cross-register the cells active in one or multiple sessions. This allows a user to track the same neurons across recording sessions. We used CellReg to register cells across sessions^26^. *Timing: 30-60 minutes*.
  - Users may alternatively use other calcium imaging cross-registration pipelines (e.g., SCOUT^27^, CaliAli^28^, etc.).
9. **Sort chemotagged cell activity**: To identify which neurons were chemogenetically activated by the CNO administration, take the cell activities present during the Chemotag session (Figure 2) and sort them based on how active each cell was. To do this, take each cell’s activity and normalize it to span between [0,1] 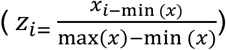. We observed that CNO began to exert its effect on neuronal activity about 5-10 minutes after the beginning of the calcium recording, during which a subpopulation of cells became highly active and others became highly inactive. Thus, we focused our analysis on minutes 10-40 of the Chemotag session. Compute the number of prominent peaks that each cell exhibits during this period. We used the *find_peaks* function using the Python package Scipy^29^. Find the optimal parameters that accurately capture the prominent peaks for the given dataset. We used the following empirically-defined parameters in the peak-finding function: height = 1e-5, distance = 7, and prominence = 0.25 (assuming a 30Hz sampling rate recording). Sort the cells based on the number of peaks they exhibited during this period (Figure 3a). Note that sorting can be based on other metrics for quantification of calcium activity, such as skew or periodicity (Figure 3b 3c). We used the number of prominent peaks as the readout to identify DREADD-activated neurons because we observed that putative DREADD-positive neurons increased their rate of calcium transients after CNO administration.
10. **Troubleshooting:** The calcium activity of each cell can be quantified by several metrics. We separately quantified two other measures–namely, skew of the area under the curve (AUC) and periodicity of oscillatory activity (Figure 3 non-GAD+ example cell shown in blue and GAD+ example cell shown in red). We computed skew because our recorded neurons typically exhibit sparse activity which produces a highly skewed AUC; however, chemogenetically activated neurons which were much more highly active displayed more normally distributed activity (Figure 3b). We computed periodicity because we observed that chemogenetically activated neurons frequently displayed periodic peaks in calcium activity, whereas endogenous neuronal activity did not exhibit periodicity (Figure 3c). This is consistent with past reports that activation of Gq-GPCRs leads to intracellular calcium oscillations, driven by endoplasmic reticulum-dependent release of calcium^18^. Decide which metric is most effective for sorting the calcium activity of the recorded cells in a given experiment to identify the neurons that are highly responsive to CNO.
11. **Assign cell identities**: Take the sorted cells from Step 9 and apply a threshold to identify the cells that are chemogenetically activated by CNO (these are the putative hM3Dq-positive neurons, representing the cell type of interest). The threshold at which neurons are categorized as chemotagged will be experiment-dependent. The expected proportions of the cell type of interest among the recorded population can be guided by the literature or from histological quantifications (see Step 12). We expected that inhibitory neurons would comprise ∼10% of all neurons in the pyramidal layer of hippocampal CA1^30^, so we assigned the top 10% of active cells as putative inhibitory neurons, while denoting the rest of the population as putative non-inhibitory neurons.
  - Notably, the threshold defined in this step can be empirically supported by testing the readout of interest using different thresholds. For example, one could compute the readout of interest assuming that the cell type of interest comprises 5% of the population, then 10%, etc., up to 100%, and see if the readout is stable in the cutoffs near the expected one (see ref^17^, Extended Figure 6).
12. **Register chemotagged cells to previous sessions**: Take the chemotagged vs non-chemotagged cells in Step 10 and register them back to cells active during the calcium imaging experiment (Step 5), using the CellReg cross-registration map^26^. The neurons recorded during the experiment were originally recorded without knowledge of their cell type identity; however, with this posthoc assignment of identity, the neurons recorded during the experiment can be delineated into subpopulations of neurons defined by their cell type.
13. **Histological validation**: Sacrifice the mice and dissect their brains. Section the brains into 50-micron brain sections, label them with a cell marker (e.g., DAPI), and mount onto slides and coverslip. With a fluorescent microscope, image the brain sections and confirm the GRIN lens is placed directly above the brain region of interest and that fluorescent expression is confined to the brain region of interest. Count the number of DREADD-positive (in our case, mCherry+) cells in the brain region of interest, relative to the number of GCaMP-labeled cells (Figure 5). The fraction of DREADD-positive cells can be used to inform the threshold to use for chemotagging (Step 10).

**Figure 1.**
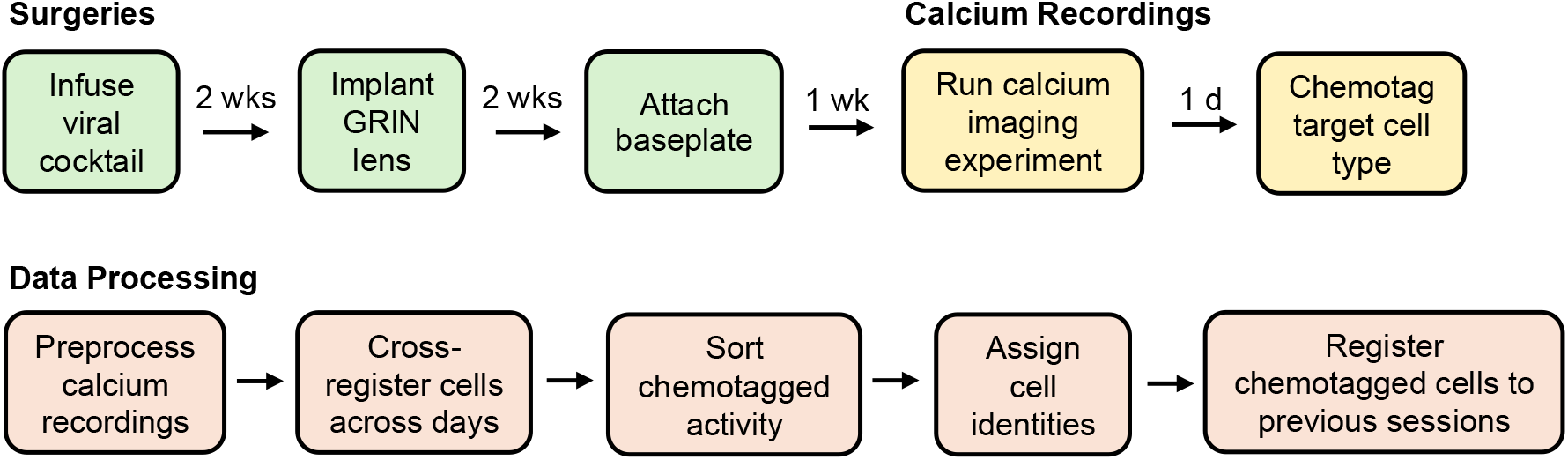
Timeline for a complete experiment using the chemotagging technique. This procedure includes performing surgeries, running a calcium imaging experiment, and processing the data thereafter for post-hoc cell-type identity assignment.

**Figure 2.**
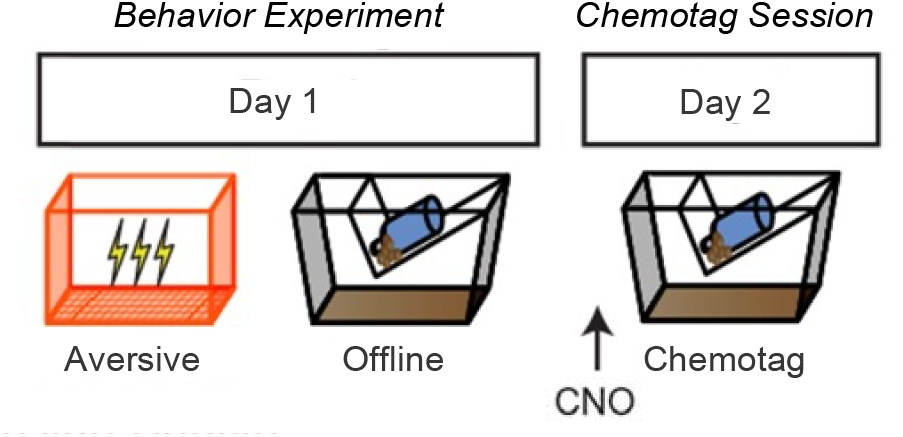
Behavioral paradigm followed by the chemotagging session.

**Figure 3.**
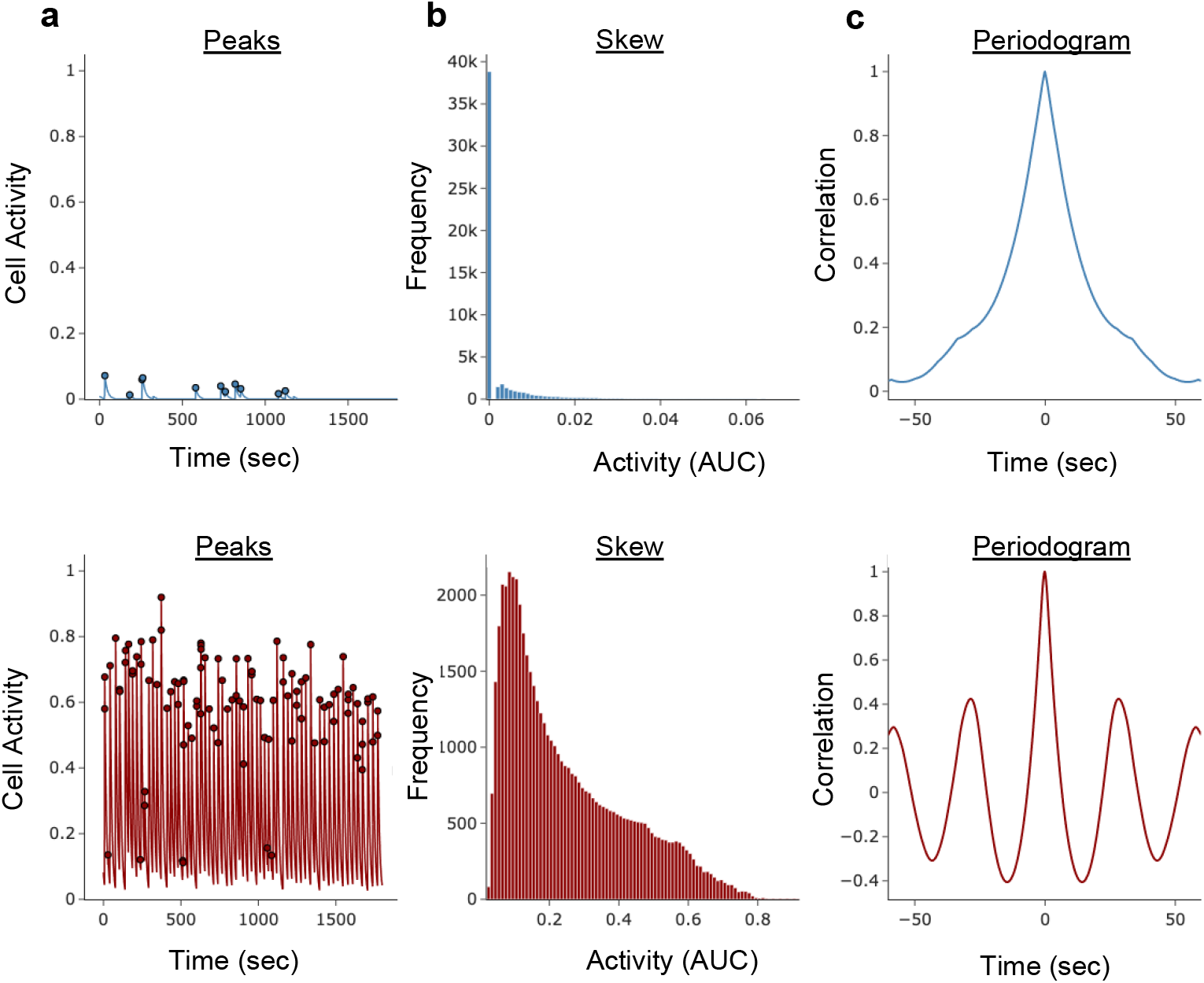
Sorting chemotagged cells with three different metrics. (a) A putative non-GAD+ cell (blue, top) and a putative-GAD+ cell (red, bottom) showing that the putative GAD+ cell displays many more peaks than the putative non-GAD+ cell. (b) Skew of the distribution of activity of the non-GAD+ cell (blue, top) and GAD+ cell (red, bottom) from Figure 4a. The sparse-firing non-GAD+ cell has a highly skewed distribution, reflecting its sparse activity, whereas the activity of the GAD+ cell is more normally distributed, reflecting its regular activity. (c) Periodogram of the non-GAD+ (blue, top) cell and GAD+ cell (red, bottom) from Figure 4a. The non-GAD+ cell exhibits non-periodic dynamics whereas the GAD+ cell displays high periodicity in its activity, reflecting its calcium oscillations.

**Figure 4.**
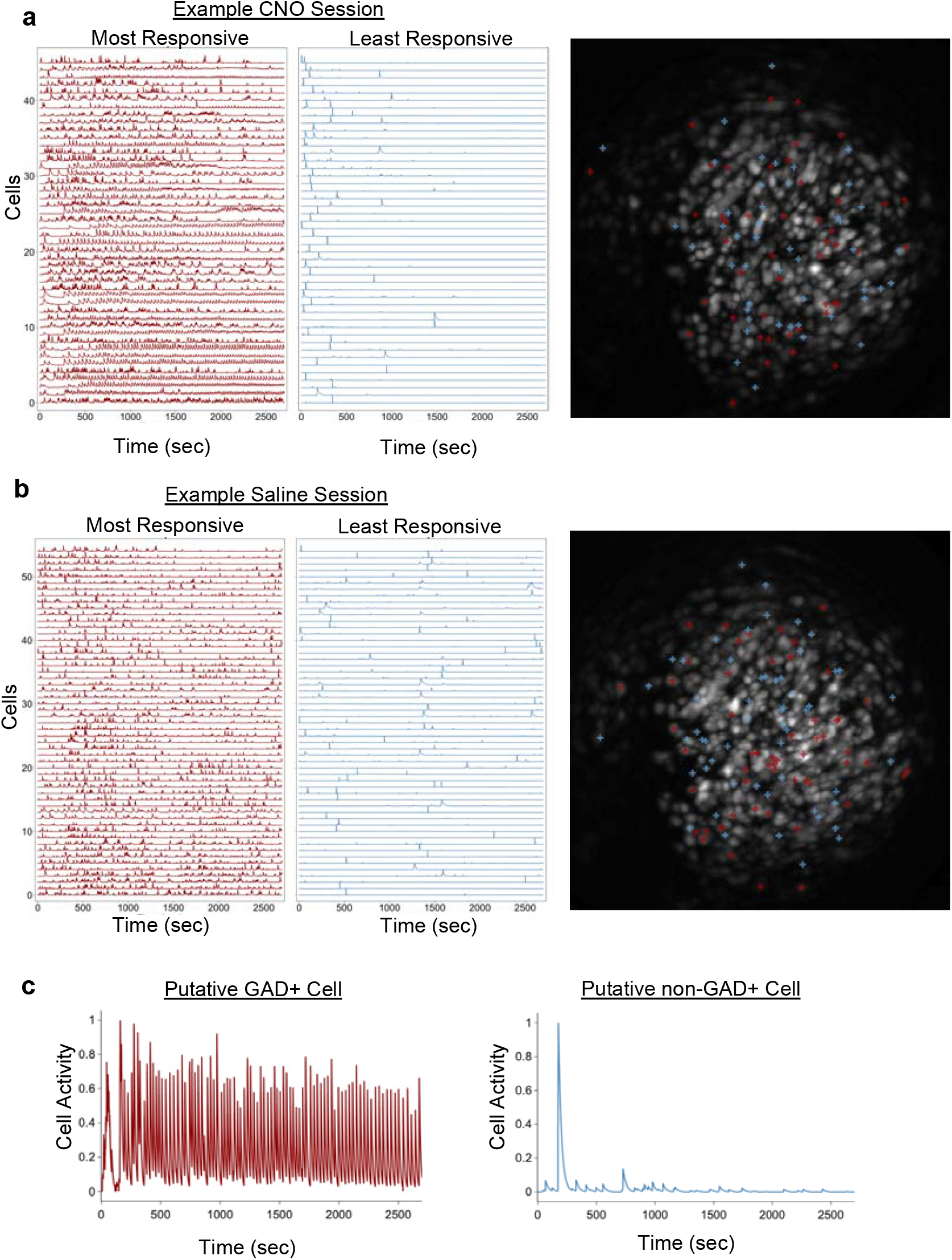
Calcium dynamics following administrations of CNO or saline. (a) 10% of most highly (red) and lowly (blue) active cells based on *in vivo* calcium activity after a representative mouse received an intraperitoneal injection of CNO. (b) Most highly (red) and lowly (blue) active cells based on *in vivo* calcium activity after mice receive an intraperitoneal injection of saline. (c) Trace of calcium activity from the chemotagging session (Figure 4a) in one representative putative GAD+ (red) and one representative putative non-GAD+ (blue) cell to highlight the delineation between typical cell activity and that of a Gq-DREADD-induced calcium oscillatory activity (adapted from ref^17^).

**Figure 5.**
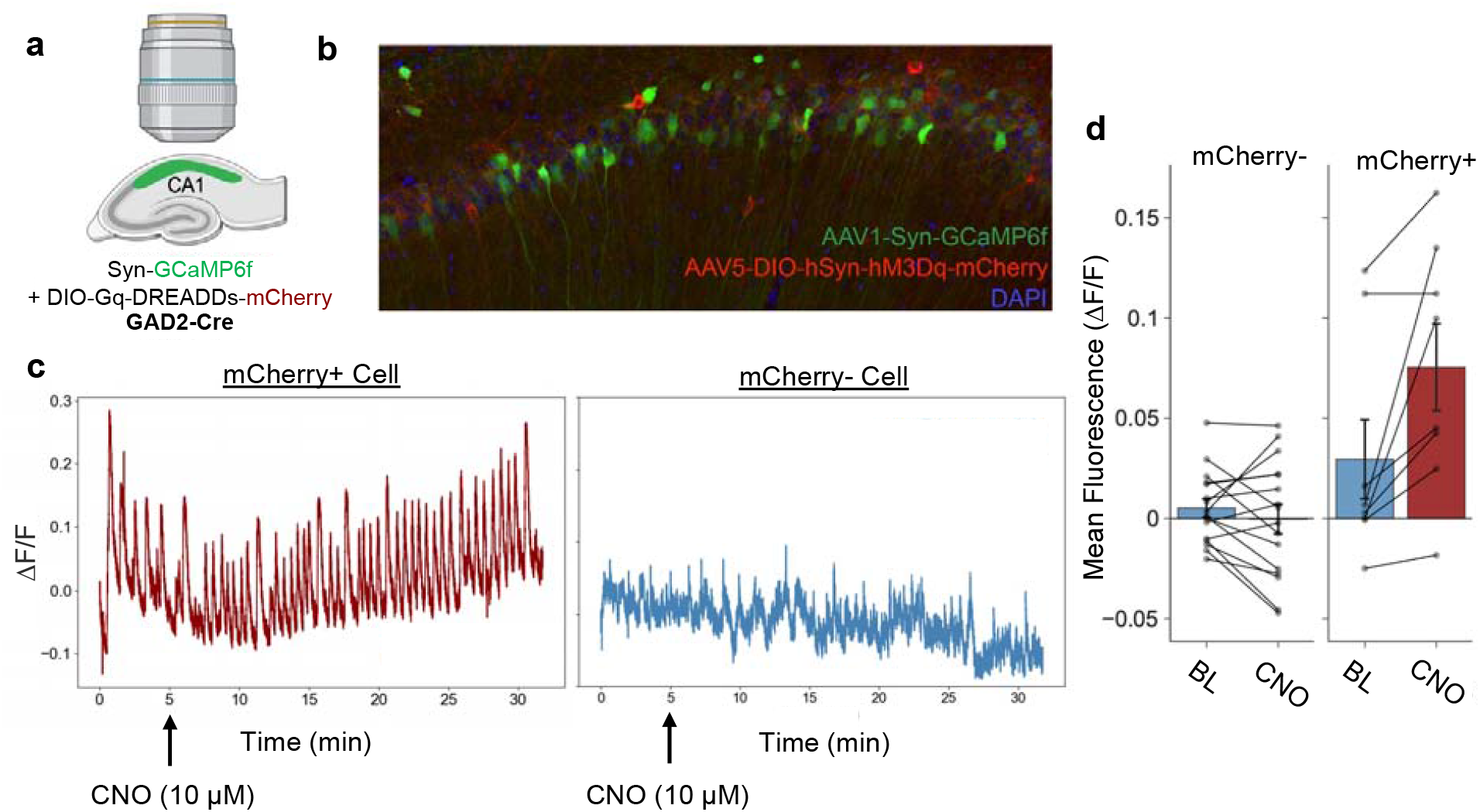
*Ex vivo* validation of CNO-induced increased calcium dynamics. (a) hippocampal slices from a Gad2-Cre mouse were prepared in vitro using a synapsin-driven GCaMP virus and a Cre-dependent excitatory DREADD virus with mCherry reporter. (b) Histological representation of GCaMP and DREADD expressing cells. (c) Calcium trace of mCherry (+) cells after CNO application (red) versus a mCherry (−) cell (blue). (d) Quantification of the change in calcium activity, as measured by fluorescence, for all cells at baseline and after CNO application. (Paired t-test with *Benjamini-Hochberg* correction; Negative: *t*_*15*_ *= 1*.*0665, p* = 0.303; mCherry: *t*_*7*_ *= 2*.*89327*, \**p* = 0.0464). Data are shown as mean ± SEM.

### Anticipated results

After the administration of CNO, a subset of cells will exhibit a robust increase in their calcium activity. Previous literature has shown that activating Gq receptors can result in distinct, robust oscillatory intracellular calcium dynamics^18-20^. Our technique leverages this known phenomenon to artificially introduce Gq receptors using a DREADD system into a cell type of interest and to selectively induce calcium oscillations within this target cell type. Because the Gq DREADD-induced calcium dynamics are distinctly oscillatory (Figure 3), the experimenter can be confident that the activity is not merely a result of the cell being highly active (i.e., firing many action potentials). Thus, cells that go into this characteristic barrage of activity upon CNO administration can be putatively labeled as being DREADD-expressing cells, and therefore, belonging to the cell type of interest. This subset of cells, the putative Gq DREADD-expressing cells, can be sorted apart from the broader population using various measures to quantify this periodic barrage of calcium activity. The putative number of DREADD-expressing cells will vary largely by the cell type of interest. Based upon the past literature of known anatomy^30^, we expected that ∼10% of cells would exhibit this behavior and thereafter be classified as DREADD-expressing cells or “chemotagged”; however, since these *in vivo* recordings do not provide ground truth about the cell type identity of each cell, a definitive threshold is difficult to determine. These thresholds should be compared with expectations from the literature, as well as from careful control experiments (e.g., histological quantifications of the fraction of cells expressing the Gq DREADD) and empirical tests to measure what fraction of cells exhibit these CNO-induced increases in periodic activity.

In a separate experiment, we conducted one chemotagging session and administered CNO to half of the mice, while the other half received saline, The following day, mice that initially received saline were injected with CNO, and mice that first received CNO were injected with saline. This allowed for a within-subjects comparison of calcium activity between mice that received an injection of saline and those same mice after CNO was introduced to their system. Once again, when mice received an injection of CNO, a subset of cells became very highly active while the rest of the population exhibited much lower activity than before the intervention (Figure 4a). Notably, this change in activity patterns was not observed when mice received a dose of saline instead (Figure 4b). Cells with a CNO-induced activity increase showed more prominent and regular activity, appearing distinctly oscillatory in the calcium trace (Figure 4c). Conversely, the remaining cell population displayed an overall decrease in their calcium dynamics. (Figure 4d).

These procedures were repeated in an *in vitro* hippocampal slice preparation, where CNO was bathed onto the slice after 5 minutes (Figure 5). Importantly, DREADD+ cells showed an increase in activity when CNO was introduced to the system, while non-DREADD+ cell activity did not change (Figure 5d).

### Limitations of chemotagging

#### Variability of CNO response

Despite the expression of the same class of DREADD receptors, variability in cellular responses to CNO can occur when using Gq-DREADD, particularly in the onset and duration of calcium oscillations. Factors contributing to this variability include differences in DREADD expression levels and receptor distribution across cells. Additionally, calcium-buffering proteins, such as parvalbumin^31^, can influence calcium dynamics, further suggesting that CNO responses may vary among different cell types.

#### Thresholds for classification

Another limitation of chemotagging is the challenge of setting appropriate thresholds for classifying cells as part of a putative subpopulation. There is currently no universally accepted metric for sorting neuronal activity, which can lead to inconsistent or arbitrary classifications. These thresholds will vary depending on the brain regions and cell types of interest. We highly recommend adjusting the criteria with careful validation of CNO responses from *ex vivo* and *in vivo* experiments.

#### Cross-registration of cells across days

One challenge in longitudinal calcium imaging experiments is the reliable cross-registration of the same neurons across multiple days of recording. This issue is exacerbated by factors such as tissue motion, variable GCaMP expression, and the potential failure of tracking cells across sessions. While registration algorithms can improve some of these challenges, further development in this area is needed to ensure reliable tracking of specific cells.

## Author contributions statements

DC, BK, MB and YZ conceived the study and designed experiments. MB and YZ conducted *in vivo* calcium imaging experiments. BK conducted *ex vivo* calcium imaging experiments. YZ and BK analyzed data. BK, MB, YZ and DC contributed to the interpretation of results and wrote the manuscript.

## Acknowledgment

This work was supported by the R01 MH120162, DP2 MH122399, R56 MH132959, Brain Research Foundation Award, Klingenstein-Simons Fellowship, NARSAD Young Investigator Award, McKnight Memory and Cognitive Disorder Award, One Mind-Otsuka Rising Star Research Award, Hirschl/Weill-Caulier Award, Friedman Brain Institute Award and Mount Sinai Distinguished Scholar Award, Chan Zuckerberg Initiative Award, McKnight Brain Research Foundation & American Foundation for Aging Research Innovator Awards in Cognitive Aging and Memory Loss Award to DJC; NIH T32HL160511 to BK; NIMH F31MH126543 to YZ. We thank all Cai lab and Shuman lab members for their comments throughout the project.

## References

1 Cardin, J. A. et al. Targeted optogenetic stimulation and recording of neurons in vivo using cell-type-specific expression of Channelrhodopsin-2. Nature protocols 5, 247–254, doi:10.1038/nprot.2009.228 (2010).

2 Roth, B. L. DREADDs for Neuroscientists. Neuron 89, 683–694, doi:10.1016/j.neuron.2016.01.040 (2016).

3 Piwecka, M., Rajewsky, N. & Rybak-Wolf, A. Single-cell and spatial transcriptomics: deciphering brain complexity in health and disease. Nature Reviews Neurology 19, 346–362, doi:10.1038/s41582-023-00809-y (2023).

4 Yao, Z. et al. A high-resolution transcriptomic and spatial atlas of cell types in the whole mouse brain. Nature 624, 317–332, doi:10.1038/s41586-023-06812-z (2023).

5 Scala, F. et al. Phenotypic variation of transcriptomic cell types in mouse motor cortex. Nature 598, 144–150, doi:10.1038/s41586-020-2907-3 (2021).

6 Gouwens, N. W. et al. Integrated Morphoelectric and Transcriptomic Classification of Cortical GABAergic Cells. Cell 183, 935-953.e919, doi:10.1016/j.cell.2020.09.057 (2020).

7 Citri, A. & Malenka, R. C. Synaptic Plasticity: Multiple Forms, Functions, and Mechanisms. Neuropsychopharmacology 33, 18–41, doi:10.1038/sj.npp.1301559 (2008).

8 Stosiek, C., Garaschuk, O., Holthoff, K. & Konnerth, A. In vivo two-photon calcium imaging of neuronal networks. Proc Natl Acad Sci U S A 100, 7319–7324, doi:10.1073/pnas.1232232100 (2003).

9 Ghosh, K. K. et al. Miniaturized integration of a fluorescence microscope. Nature Methods 8, 871–878, doi:10.1038/nmeth.1694 (2011).

10 Cai, D. J. et al. A shared neural ensemble links distinct contextual memories encoded close in time. Nature 534, 115–118, doi:10.1038/nature17955 (2016).

11 Aharoni, D. & Hoogland, T. M. Circuit Investigations With Open-Source Miniaturized Microscopes: Past, Present and Future. Frontiers in cellular neuroscience 13, 141, doi:10.3389/fncel.2019.00141 (2019).

12 Dong, Z. et al. Simultaneous dual-color calcium imaging in freely-behaving mice. bioRxiv, doi:10.1101/2024.07.03.601770 (2024).

13 Papaioannou, S. & Medini, P. Advantages, Pitfalls, and Developments of All Optical Interrogation Strategies of Microcircuits in vivo. Front Neurosci 16, 859803, doi:10.3389/fnins.2022.859803 (2022).

14 Oda, K. et al. Crystal structure of the red light-activated channelrhodopsin Chrimson. Nature Communications 9, 3949, doi:10.1038/s41467-018-06421-9 (2018).

15 Chen, T. W. et al. Ultrasensitive fluorescent proteins for imaging neuronal activity. Nature 499, 295–300, doi:10.1038/nature12354 (2013).

16 Akerboom, J. et al. Genetically encoded calcium indicators for multi-color neural activity imaging and combination with optogenetics. Front Mol Neurosci 6, 2, doi:10.3389/fnmol.2013.00002 (2013).

17 Zaki, Y. et al. Offline ensemble co-reactivation links memories across days. Nature, doi:10.1038/s41586-024-08168-4 (2024).

18 McDonough, R. C., Gilbert, R. M., Gleghorn, J. P. & Price, C. Targeted Gq-GPCR activation drives ER-dependent calcium oscillations in chondrocytes. Cell Calcium 94, 102363, doi:10.1016/j.ceca.2021.102363 (2021).

19 Alexander, G. M. et al. Remote control of neuronal activity in transgenic mice expressing evolved G protein-coupled receptors. Neuron 63, 27–39, doi:10.1016/j.neuron.2009.06.014 (2009).

20 Rogan, S. C. & Roth, B. L. Remote Control of Neuronal Signaling. Pharmacological Reviews 63, 291–315, doi:10.1124/pr.110.003020 (2011).

21 Tanaka, K. Z. et al. The hippocampal engram maps experience but not place. Science 361, 392–397, doi:10.1126/science.aat5397 (2018).

22 Resendez, S. L. et al. Visualization of cortical, subcortical and deep brain neural circuit dynamics during naturalistic mammalian behavior with head-mounted microscopes and chronically implanted lenses. Nature protocols 11, 566–597, doi:10.1038/nprot.2016.021 (2016).

23 Dong, Z. et al. Minian, an open-source miniscope analysis pipeline. eLife 11, doi:10.7554/eLife.70661 (2022).

24 Giovannucci, A. et al. CaImAn an open source tool for scalable calcium imaging data analysis. Elife 8, doi:10.7554/eLife.38173 (2019).

25 Lu, J. et al. MIN1PIPE: A Miniscope 1-Photon-Based Calcium Imaging Signal Extraction Pipeline. Cell Rep 23, 3673–3684, doi:10.1016/j.celrep.2018.05.062 (2018).

26 Sheintuch, L. et al. Tracking the Same Neurons across Multiple Days in Ca(2+) Imaging Data. Cell reports 21, 1102–1115, doi:10.1016/j.celrep.2017.10.013 (2017).

27 Johnston, K. G. et al. Tracking longitudinal population dynamics of single neuronal calcium signal using SCOUT. Cell Rep Methods 2, 100207, doi:10.1016/j.crmeth.2022.100207 (2022).

28 Vergara, P. et al. CaliAli, a tool for long-term tracking of neuronal population dynamics in calcium imaging. bioRxiv, 2023.2005.2019.540935, doi:10.1101/2023.05.19.540935 (2023).

29 Virtanen, P. et al. SciPy 1.0: fundamental algorithms for scientific computing in Python. Nat Methods 17, 261–272, doi:10.1038/s41592-019-0686-2 (2020).

30 Bezaire, M. J. & Soltesz, I. Quantitative assessment of CA1 local circuits: knowledge base for interneuron-pyramidal cell connectivity. Hippocampus 23, 751–785, doi:10.1002/hipo.22141 (2013).

31 Haiech, J., Moreau, M., Leclerc, C. & Kilhoffer, M. C. Facts and conjectures on calmodulin and its cousin proteins, parvalbumin and troponin C. Biochim Biophys Acta Mol Cell Res 1866, 1046–1053, doi:10.1016/j.bbamcr.2019.01.014 (2019).

